# Cholinergic interneurons of the dorsomedial striatum mediate winner-loser effects on social hierarchy dynamics in male mice

**DOI:** 10.1101/2024.12.04.626719

**Authors:** Mao-Ting Hsu, Yumiko Akamine, Kiyoto Kurima, Kazumasa Z. Tanaka, Jeffery R. Wickens

## Abstract

Cholinergic interneurons of the dorsomedial striatum may play a role in social hierarchy dynamics. A social hierarchy is an organization of individuals by rank that occurs in social animals. Establishing a new social hierarchy involves flexible behavior in deciding whether to be a winner or loser, experience of winning or losing, and stabilization of rank. The neural circuits underlying such flexible behavior have yet to be fully understood, but previous research indicates that cholinergic interneurons in the dorsomedial striatum play a role in behavioral flexibility. We used the dominance tube test to measure ranking within group housed mice, before and after between-cage competitions using the same test. We found that the experience of winning or losing against mice from different cages not only contributes to new social hierarchies among the competitors, but also causally influences the subsequent social hierarchy among their cage mates in the home cage – supporting the hypothesis of winner-loser effects on later social ranking. To test the hypothesis that cholinergic interneurons contribute to social hierarchy dynamics, we made a selective lesion of cholinergic interneurons in the dorsomedial striatum. The lesion did not prevent social hierarchy formation among pairs of similarly ranked individuals from different cages. However, it reduced the loser effect of external competition on the subsequent home-cage rankings in dominant mice. In light of these results we suggest that cholinergic interneurons in dorsomedial striatum increase the flexibility of social hierarchy dynamics.

**Significance statement:** The effect of winning or losing a competition on subsequent ranking in mouse home cage social hierarchies was examined using the dominance tube test. We found that losing, when dominant mice were defeated by equally ranked mice from another cage, led to decreased social rank in their home cage. Conversely, winning by initially subordinate mice led to increased rank in the home cage social hierarchy. The loser effect on subsequent behavior in dominant mice was reduced after selective lesions of the cholinergic interneurons of the dorsomedial striatum. We suggest that losing might produce these effects by altering the activity of cholinergic interneurons, and thus modulating synaptic plasticity in neural circuits involved in flexible decision making and positive reinforcement.

## Introduction

Social hierarchy is the ranking of members of a social group, and occurs widely in social animals, such as rodents, non-human primates, and humans (1–3). A stable social hierarchy has the adaptive advantage of efficiently distributing resources to individuals according to their rank, without repeated challenges. A dominant individual exercises dominance in multiple aspects of behavior including territory defense, food access priority, and mating priority. Conversely, a subordinate shows submissiveness to a dominant, and this submissiveness prevents needless competition in a social group (3). Several authors have proposed that the reinforcing effects of experiencing winning or losing are important for social hierarchy formation (4–7). This “winner-loser effect” hypothesis states that winners become more likely to escalate a contest, while losers become less willing to engage in conflicts (8). For example, male swordtails with prior winning experience tend to occupy a dominant position in the newly grouped society, whereas individuals with prior losing experience tend to take a subordinate position (9). Unexpected winning or losing also seems to reshape the social hierarchy within the same cage of mice (10, 11). These findings have led to the suggestion that the winner-loser effect may induce hierarchical flexibility within a social group. Recent research indicates that the medial prefrontal cortex regulates the effortful dominance behaviors in mice (11, 12). However, the knowledge of the neural mechanisms of social hierarchy formation is still limited.

The dorsomedial striatum (DMS), one of the downstream pathways of the medial prefrontal cortex, integrates inputs from cortex, thalamus, midbrain, and limbic systems. The behavioral functions of DMS include behavioral flexibility and goal-directed behavior (13–15). In relation to social hierarchy, behavioral flexibility may provide a useful conceptual framework for understanding switching between being a winner or loser. Thus, DMS may be involved in hierarchical flexibility and social hierarchy formation. Among different types of neurons in a striatum, cholinergic interneurons (ChIs) represent as few as 1% of the striatal neurons in rodents (16), but they are large and have widely arborized dendrites and axons (17). These interneurons have been reported to regulate goal directed behavior, behavioral flexibility, and set-shifting (18–22). Thus, we hypothesize that ChIs in DMS regulate the hierarchical flexibility and play a critical role in social hierarchy dynamics.

We investigated social hierarchy formation and maintenance in male mice focusing on strategy switching after competition in the dominance tube test (DTT). We then tested the hypothesis that DMS ChIs play a causal role in these winner-loser effects. A sequence of pairwise competitions between mice of equal social rank was used to test how winning or losing affected the social hierarchy in the home cages. Selective lesions of ChIs in DMS were used to determine their involvement in the effects of winning or losing.

## Results

### Effect of external winning or losing experience on a new social hierarchy formation

The behavioral procedure is outline in Figure 1. Social ranks within home cages were determined by repeated dominance tube tests (DTTs) of all possible pairings of mice from each cage over 4-6 days. In this phase a total of 114 mice were examined, grouped into 38 home-cage groups of 3 mice in each cage. These mice were divided into two groups in this behavioral procedure: naïve (69 mice, 23 home-cage groups, experienced between-cage competition) and control (45 mice,15 home-cage groups, did not experience between-cage competition). A transitive hierarchy was considered to exist when individual mice held unique dominance ranks (dominant, intermediate, or subordinate) that were stable over four consecutive days or five out of six days – Criterion I – based on a previous study (11). No mice or cage groups were excluded from this analysis and C57BL/6J and ChAT-Cre Het mice were pooled after determining there was no significant difference in prevalence of transitive hierarchy in these strains (*p* = 0.2251, 2×2 Fisher’s exact test). Initially, a transitive hierarchy was observed in 28 of the 38 cages, a prevalence of about 73% (Table 1). To increase the sample size of series sequence of pairwise competitions between mice of equal social, we defined a different criterion – Criterion II – for the cages which showed a non-transitive hierarchy under Criterion I. The mouse that won most often in the initial within-cage DTT was designated dominant. The mouse that won least often was designated subordinate. The mouse was designated intermediate if its winning rate was between the dominant and subordinate. If the winning rates were the same, we defined the mouse status according to their hierarchical status in the last session.

**Figure 1.**
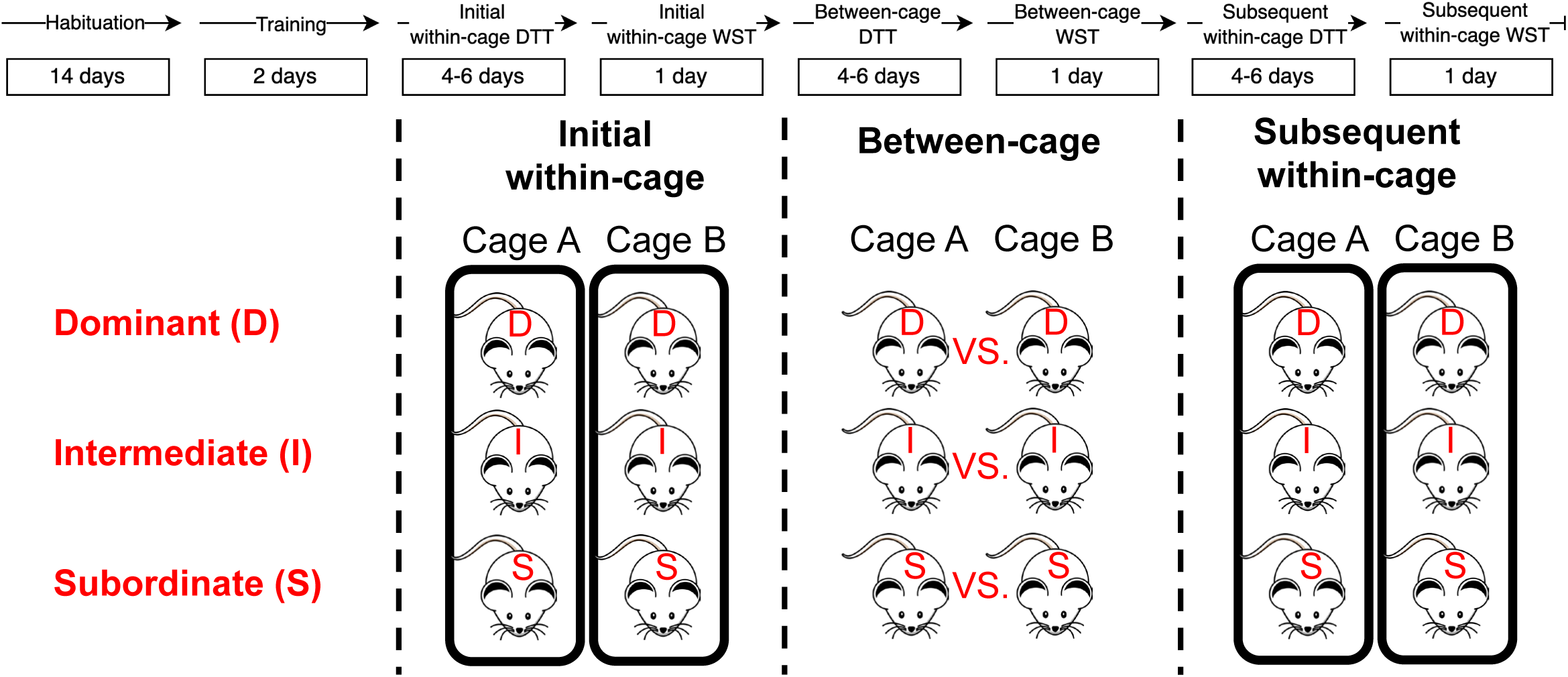
The behavioral procedure of series of DTT and WST. The initial within-cage is to recognize the initial social hierarchy in each home cage of mice. The between-cage is to establish new social hierarchies between mice, positioned in the same ranking, from different cages through pairwise competition of DTT. The subsequent within-cage is to investigate the effect of the external winning or losing experience on the social hierarchy in the home cages.

**Table 1.** Initial prevalence of transitive hierarchy in male mice.

| Type of Social Hierarchy | C57BL/6J (cages) | ChAT-Cre (cages) | Total cages |
| --- | --- | --- | --- |
| Transitive | 10 | 18 | 28 |
| Non-Transitive | 1 | 9 | 10 |

There is no significant difference between C57BL/6J and ChAT-Cre mice ( $p = 0.2251$ , Fisher's exact test).

To test whether mice displayed the winner-loser effect (8), mice of each social rank from different cages competed with mice of the same rank from different cages (dominant vs. dominant, intermediate vs. intermediate, and subordinate vs. subordinate), in repeated between-cage DTTs. The duration of the competition in the first and last session was measured (Figure 2A). A two-way repeated measures ANOVA was applied to analyze these numerical data. The statistical results showed that there was a significant mean difference between the first and last session (F_1,8_ = 19.4853, *p* = 0.0022) and between social ranks (F_2,16_ = 28.5554, *p* < 0.001). There was also an interaction between the session and social ranks (F_2,16_ = 24.1155, *p* < 0.001). In the multiple comparison analysis, there was a significant decrease in duration from the first to last session in competition between home-cage dominant mice from different cages (*p* < 0.001, Tukey’s HSD). This indicates that a new social hierarchy was established among dominants after experiencing a win or loss.

**Figure 2.**
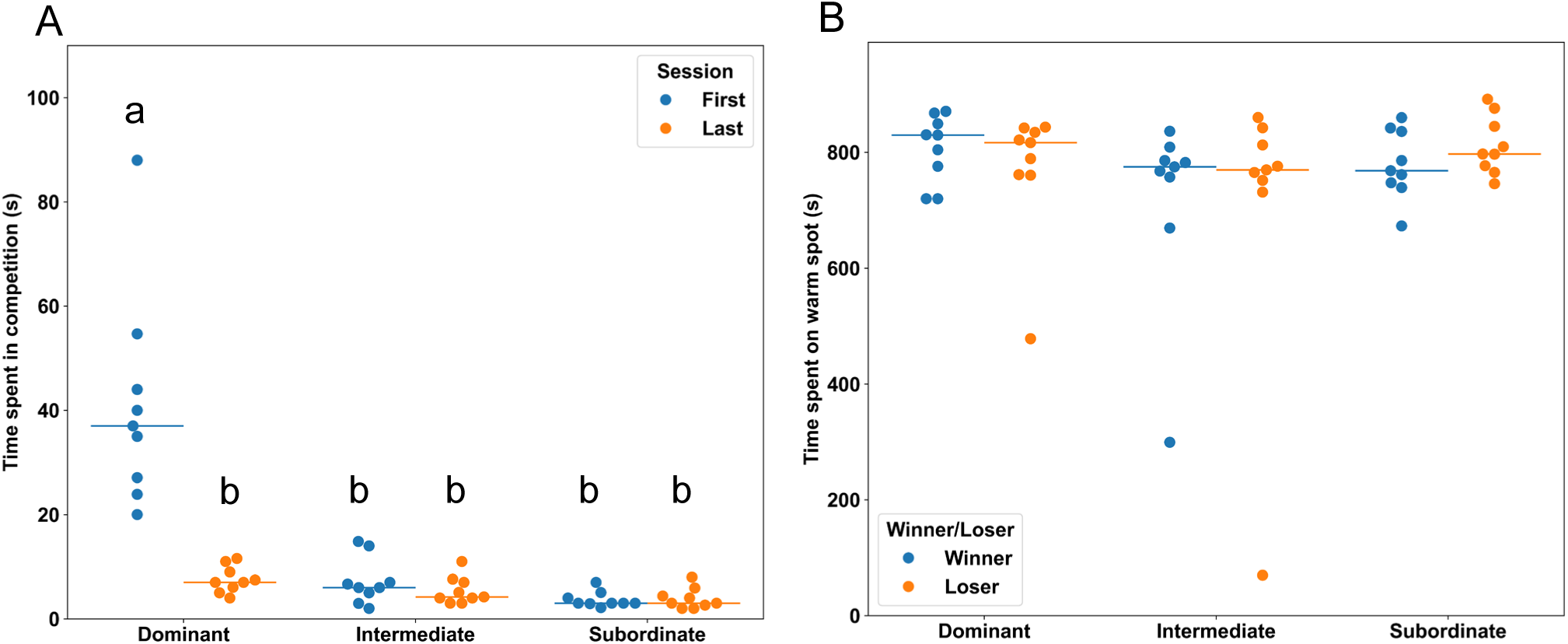
(**A**) Time spent in the competition by rank in the between-cage DTT. Blue dots indicate individual competition time in the first session. Orange dots indicate individual competition time in the last session. Line shows mean of first or last session for each rank. (**B**) Time spend on the warm spot by rank in the between-cage WST. Blue dots indicate individual time winners in the between-cage DTT spent on the warm spot. Orange dots indicate the individual time losers in the between-cage DTT spent on warm spot. Line shows mean of the time spent on the warm spot of the winners or losers in the between-cage DTT in each rank.

In contrast to the initially prolonged competition among dominant-dominant pairs, competition duration was initially short in the intermediate-intermediate and subordinate-subordinate competitions, and, over sessions, there was no significant change in the duration of the competition in these pairings (intermediate-intermediate, *p* = 0.9981; subordinate-subordinate, *p* = 1, Tukey’s HSD). This suggests that among the intermediates and subordinates the prior experience of losing causes one or other of them to quickly retreat in the DTT. In addition, this result also showed a coherence to a previous study which indicated that lower-ranked male mice presented higher susceptible to chronic social defeat stress task than dominants (23). The intermediates and subordinates may quickly decide to withdraw from competitions due to their high susceptibility. Noteworthy, all competitions among mice of each social rank remained stable for four consecutive days. This result suggested that a new social hierarchy was established among dominants, intermediates, and subordinates after experiencing a win or loss.

Mice were also tested on the WST after each DTT. However, when winners in the between-cage DTT were compared with losers on the WST, there was no significant difference (Figure 2B) (Dominant: *p* = 0.4181; Intermediate: *p* = 0.9078; Subordinate: *p* = 0.2274, Student’s *t*-test). As shown in supplementary figure 1, there was no correlation (initial and subsequent within-cage) between the rank determined by DTT and dominance performance in WST on the initial and subsequent within-cage competition (initial within-cage (Supplementary figure 1C) – *p* = 0.8326 and R^2^ = 0.0009; subsequent within-cage (Supplementary figure 1D) – *p* = 0.3665 and R^2^ = 0.0157, Pearson’s correlation test). This may be due to the relatively small size of each cage group reducing the competition for the warm spot test.

Between-cage competition may affect the dominance performance in WST. Therefore, we compared dominance performance between DTT and WST in the control group which showed a stable hierarchy between initial and subsequent within-cage DTT. As shown in supplementary figure 2, there was no correlation (initial and subsequent within-cage) between the rank determined by DTT and dominance performance in WST on the initial and subsequent within-cage competition (initial within-cage (Supplementary figure 2D) – *p* = 0.899 and R^2^ = 0.0007; subsequent within-cage (Supplementary figure 2E) – *p* = 0.6615 and R^2^ = 0.0078, Pearson’s correlation test). This indicated that the between-cage competition may not affect the dominance performance in WST. On the other hand, the rank determined by initial WST was correlated with dominance performance in subsequent WST (Supplementary figure 2F) – *p* < 0.001 and R^2^ = 0.6677, showing there was consistent dominance performance over time in WST. In the multiple comparison, there was significant difference in dominant-subordinate (*p* < 0.001, Tukey’s HSD) and intermediate-subordinate (*p* < 0.001, Tukey’s HSD) comparison. However, there was no significant difference in the dominant-intermediate (*p* = 0.1985, Tukey’s HSD) comparison. This indicated that competition in WST may escalate between the higher ranked mice in the relatively small size of each cage group.

### Effect of external winning or losing experience on social ranking

The between-cage DTT caused changes in the social hierarchy in the home cages, as determined by subsequent within-cage DTTs (Table 2). The social ranking of the nine dominant mice who won in the between-cage DTT did not change (*p* = 1 compared with the control, 2×2 Fisher’s exact test, Table 2A). However, six out of the nine dominant mice who lost in the between-cage DTT subsequently lost to lower ranked mice in the home-cage DTT, thus decreasing in social rank. This indicates a significant effect of the between-cage competition outcome on the outcome of the subsequent within-cage competition (*p* = 0.0037 compared with the control, 2×2 Fisher’s exact test). There was also a significant difference between dominant-winner and dominant-loser (*p* = 0.009, 2×2 Fisher’s exact test). These results are consistent with the winner-loser effect, in that dominant mice that experienced winning in the between-cage DTT were more likely to win in the subsequent within-cage DTT, while dominant mice that experienced losing in the between-cage DTT were more likely to lose in the subsequent within-cage DTT.

**Table 2.**
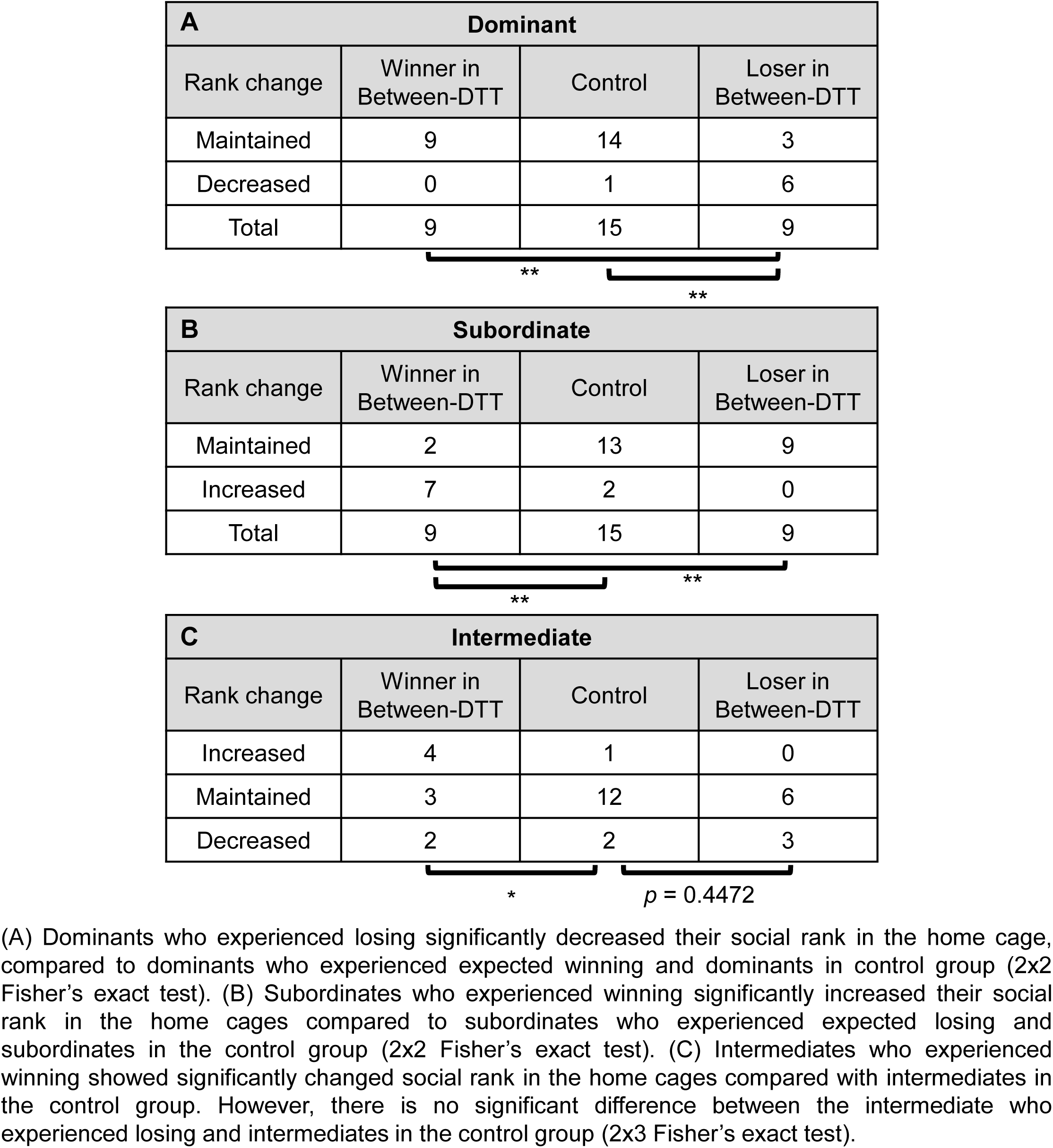
Rank change in the home cages after the between-cage DTT competition.

| **A Dominant** |  |  |  |
| --- | --- | --- | --- |
| Rank change | Winner in Between-DTT | Control | Loser in Between-DTT |
| Maintained | 9 | 14 | 3 |
| Decreased | 0 | 1 | 6 |
| Total | 9 | 15 | 9 |
| **<br>** |  |  |  |

| **B Subordinate** |  |  |  |
| --- | --- | --- | --- |
| Rank change | Winner in Between-DTT | Control | Loser in Between-DTT |
| Maintained | 2 | 13 | 9 |
| Increased | 7 | 2 | 0 |
| Total | 9 | 15 | 9 |
| **<br>** |  |  |  |

| **C Intermediate** |  |  |  |
| --- | --- | --- | --- |
| Rank change | Winner in Between-DTT | Control | Loser in Between-DTT |
| Increased | 4 | 1 | 0 |
| Maintained | 3 | 12 | 6 |
| Decreased | 2 | 2 | 3 |
| *<br>$p = 0.4472$ |  |  |  |

(A) Dominants who experienced losing significantly decreased their social rank in the home cage, compared to dominants who experienced expected winning and dominants in control group (2x2 Fisher's exact test). (B) Subordinates who experienced winning significantly increased their social rank in the home cages compared to subordinates who experienced expected losing and subordinates in the control group (2x2 Fisher's exact test). (C) Intermediates who experienced winning showed significantly changed social rank in the home cages compared with intermediates in the control group. However, there is no significant difference between the intermediate who experienced losing and intermediates in the control group (2x3 Fisher's exact test).

The converse effect was observed in the subordinate mice after competition with mice from other cages (Table 2B). Most of the initially subordinate mice that won in the between-cage competition increased (7/9) their status in subsequent within-cage competition (*p* = 0.0029 compared with the control, 2×2 Fisher’s exact test). On the other hand, 9/9 subordinate mice that lost in the between-cage competition did not increase their status (*p* = 0.5108 compared with the control, 2×2 Fisher’s exact test). There was also a significant difference between subordinate-loser and subordinate-winner (*p* = 0.0022, 2×2 Fisher’s exact test).

Intermediate ranked mice who won or lost in the between-cage DTT also showed changes in the subsequent home-cage competitions, but given that there were three possible outcomes, the effects are more complex (Table 2C). Intermediates who won in the between-cage DTT increased, maintained, or decreased their status, resulting in a significant difference from the control (*p* = 0.03, 2×3 Fisher’s exact test). Intermediates who lost in the between-cage DTT showed no significant differences when compared with the control (*p* = 0.4472, 2×3 Fisher’s exact test).

### Effect of external winning or losing experience of cage-mates on social ranking

The experience of cage-mates in the between-cage competitions also appeared to affect the outcome of subsequent home-cage competitions over days (Figure 3). All dominant rank mice who won in the between-cage DTT tended to maintain their social rank in subsequent within-cage DTT (9/9). On the other hand, six out of seven of the previously dominant mice who subsequently lost in the home-cage DTT (decreased their status) were competing against lower-ranked cage-mates that had won in the between-cage DTT. This loser effect reduced their ranks compared with the control dominants (day1 – *p* < 0.001, day2 – *p* < 0.001, day3 – *p* = 0.009, day4 – *p* < 0.001, Dunn’s test, Bonferroni adjustment) and the dominant mice who won the competition in between-cage DTT (day1 – *p* < 0.001, day2 – *p* < 0.001, day3 – *p* = 0.003, day4 – *p* < 0.001, Dunn’s test, Bonferroni adjustment) over four consecutive days (Figure 3A). Dominants who lost in the between-cage DTT maintained their dominant ranks in the context that their cage-mates also lost in the between-cage DTT (2/2).

**Figure 3.**
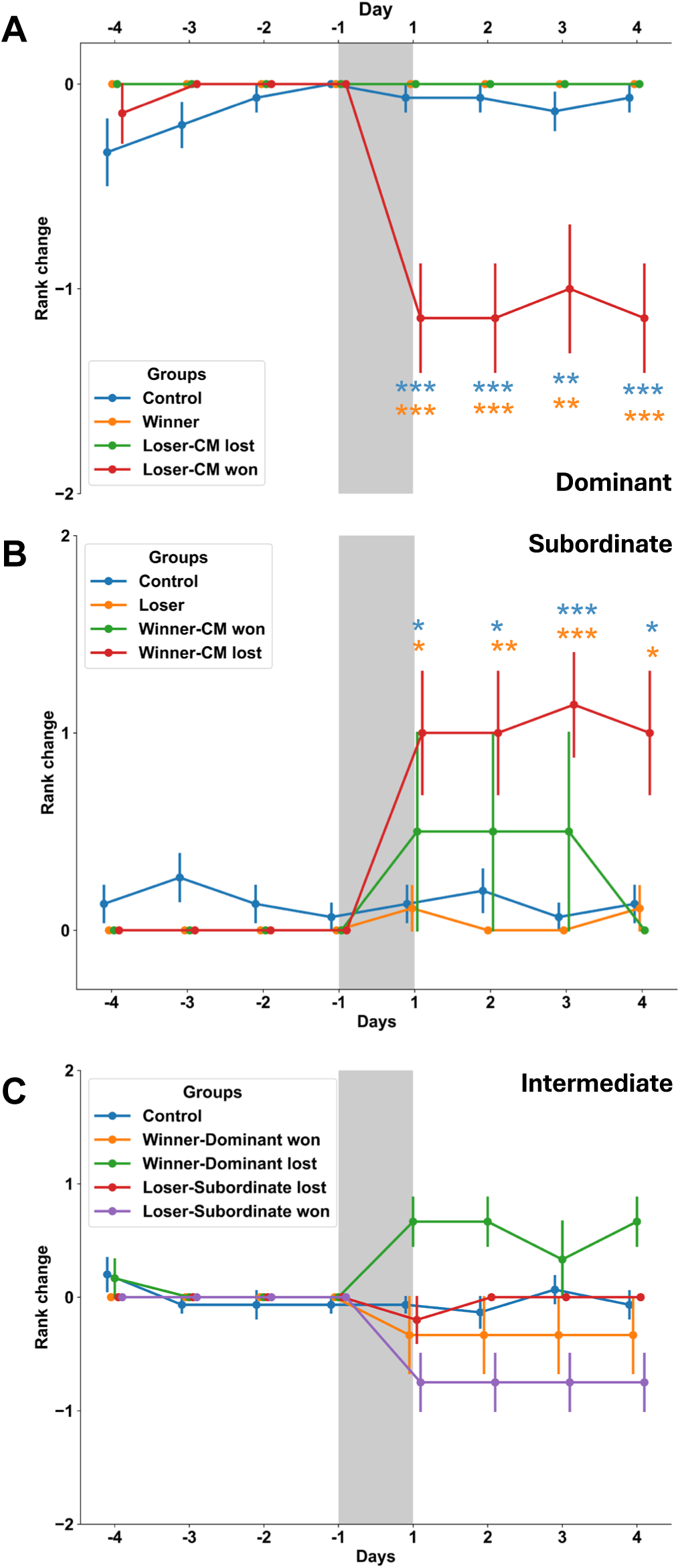
Rank change of subsequent hierarchy after experiencing between-cage competition. Grey area represents phase of between-cage competition. Horizontal axis indicates the dates before and after between-cage competition. (**A**) Dominants who experienced losing in the context that one or two of their cage-mates won in the between-DTT significantly decreased their ranks in home-cages over four consecutive days (n = 7) compared with dominants who experienced winning (orange asterisks) or dominants in control group (blue asterisks). (**B**) Subordinates who experienced winning in the context that one or two of their cage-mates lost in the between-DTT significantly increased their ranks in home-cages over four consecutive days (n = 7) compared with subordinates who experienced losing (orange asterisks) or subordinates in control group (blue asterisks). (**C**) There is no significant difference between intermediates who experienced winning in the context that their dominants lost in the between-DTT and intermediates in the control group over four consecutive days (n = 6). Moreover, there is no significant difference between intermediates who experienced losing in the context that their subordinate won in the between-DTT and intermediates in the control group over four consecutive days (n = 4) (Dunn’s test, Bonferroni adjustment). (Cage-mates: CM)

The winner-loser effect was also evident among the mice with subordinate rank in their home cage (Figure 3B). All subordinate rank mice who lost in the between-cage DTT tended to maintain their social rank in subsequent within-cage DTT (9/9). However, the winners of competitions between subordinate rank mice in the between-cage DTT tended to increase their social rank when competing against a higher-ranking mouse who lost in the between-cage DTT (six out of seven mice). This winner effect elevated their ranks compared with the control subordinates (day1 – *p* = 0.0169, day2 – *p* = 0.0433, day3 – *p* < 0.001, day4 – *p* = 0.013, Dunn’s test, Bonferroni adjustment) and the subordinate mice who lost the competition in between-cage DTT (day1 – *p* = 0.03, day2 – *p* = 0.0067, day3 – *p* < 0.001, day4 – *p* = 0.0239, Dunn’s test, Bonferroni adjustment) over four consecutive days. Conversely, if all cage mates won in the between-cage DTT then the subordinate mice tended to maintain their status whether they won or lost in the between-cage DTT (2/3).

In the case in which intermediates who won the between-cage DTT had dominant cage-mates that lost in the between-cage DTT (n = 6), there were no significant differences from control intermediate (day1 – *p* = 0.0833, day2 – *p* = 0.079, day3 – *p* = 1, day4 – *p* = 0.1047, Dunn’s test, Bonferroni adjustment) over four consecutive days (Figure 3C). In the case in which intermediates who lost the between-cage DTT had subordinate cage-mates that won in the between-cage DTT (n = 4), there were no significant differences from control intermediate (day1 – *p* = 0.2436, day2 – *p* = 0.6536, day3 – *p* = 0.1221, day4 – *p* = 0.3557, Dunn’s test, Bonferroni adjustment).

Taken together, the foregoing results indicate an association between the experience of winning or losing experience in the between-cage DTT, and the outcome of subsequent within-cage DTTs. The prior experience of both mice in the competition appears to be important.

### Lesion of cholinergic interneurons did not reduce the prevalence of transitive hierarchy

The winner-loser effect is a form of behavioral flexibility in which the outcome of a competition is followed by a change in strategy. Previous work has shown that striatal cholinergic interneurons are involved in behavioral flexibility (18, 21). To investigate the role of cholinergic interneurons in the effects observed, we selectively ablated cholinergic interneurons using the Cre-dependent diphtheria AAV (AAV1-mCherry-flex-dtA) in ChAT-Cre mice. In these experiments a total of 48 ChAT-cre transgenic animals were injected with virus. Fluorescent immunostaining showed that cholinergic interneurons in DMS were selectively ablated while non-cholinergic cells were preserved (Figure 4). Lesioning the DMS cholinergic interneurons did not cause a significant difference in the initial prevalence of transitive hierarchies, which determined by Criterion I, in the home cages prior to experimental competition sessions (unlesioned 28/38 vs lesioned 8/16, *p* = 0.1193, 2×2 Fisher’s exact test, Table 3). These findings indicate that the ChI lesion did not change the prevalence of transitive hierarchy.

**Figure 4.**
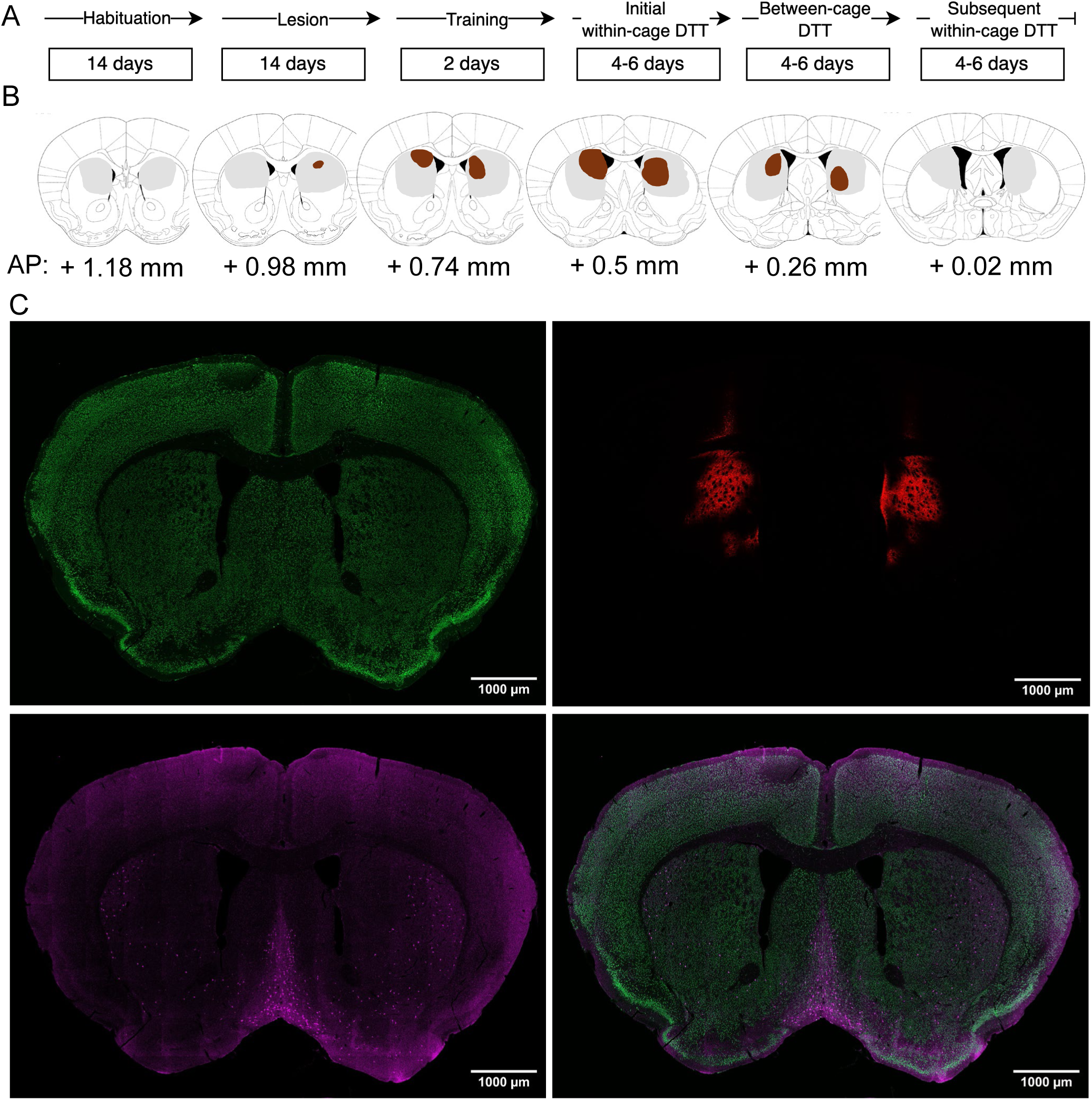
The behavioral procedure and histological validation of the ablation of the cholinergic interneurons in the DMS. (A) Experimental procedure involving striatal cholinergic interneuron ablation and dominance tube test. (B) The maximum (grey) and minimum (brown) areas of expression of the virus in DMS among 48 mice. (C) Green represents NeuN expression. Red shows the mCherry expression. Magenta demonstrates the ChAT expression. Bottom right shows merge of NeuN and ChAT imaging. The scale bar is 1 mm.

**Table 3.** Prevalence of the transitive hierarchy in male mice in the initial within-cage DTT.

| Type of Social Hierarchy | Naïve | Lesion |
| --- | --- | --- |
| Transitive | 28 | 8 |
| Non-Transitive | 10 | 8 |

There is no significant difference between Naïve and Chl Lesion group. (Unit: Cage of mice) ( $p = 0.1193$ , 2x2 Fisher's exact test)

### Lesion of cholinergic interneurons reduces the loser effect on dominant mice

In ChI-lesioned mice (Figure 5), as in unlesioned mice, we used two-way repeated measures ANOVA to analyze the numerical data. The statistical results showed that there was a significant mean difference between the first and last session (F_1,7_ = 24.9865, *p* = 0.0015) and between social ranks (F_2,14_ = 31.1716, *p* < 0.001). There was also an interaction between the session and social ranks (F_2,14_ = 13.3862, *p* < 0.001). In the multiple comparison analysis, the duration of dominant-dominant competition was significantly reduced (*p* < 0.001, Tukey’s HSD). Also in lesioned mice, as previously in unlesioned mice, the duration of competition was initially short in intermediate-intermediate (*p* = 0.9288, Tukey’s HSD) and subordinate-subordinate (*p* = 0.9955, Tukey’s HSD) competition and there was no significant reduction in duration. This result indicates that the lesion of ChIs in DMS does not alter new social hierarchy formation after competition between two individuals with the same social rank.

**Figure 5.**
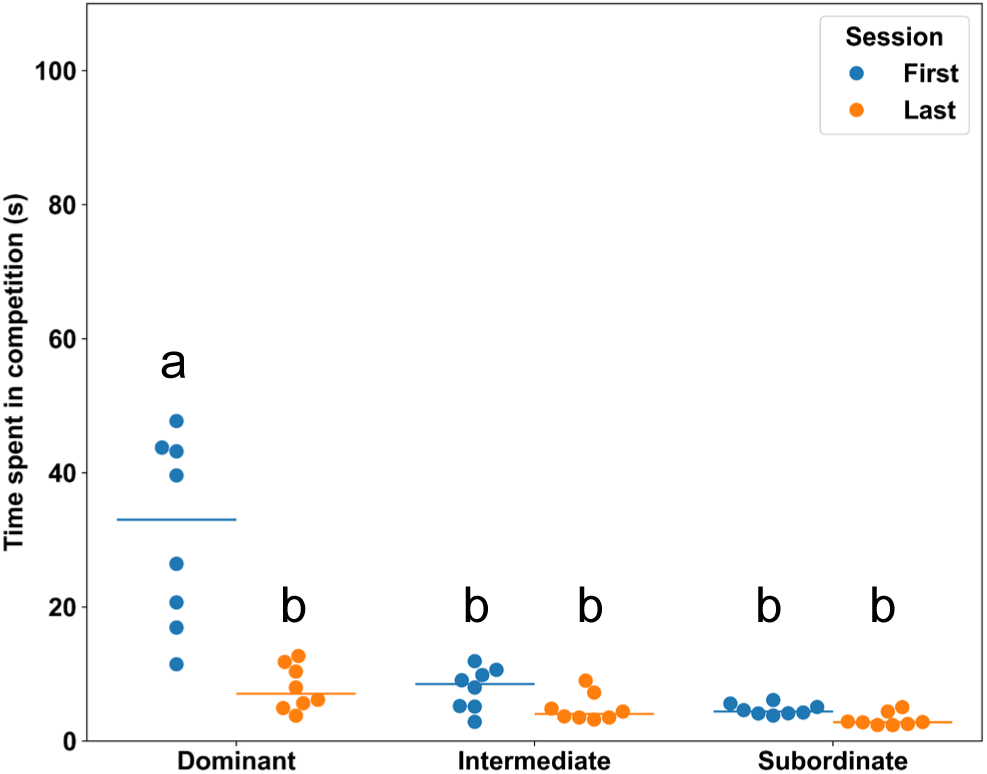
Time spent in competition by social rank in the between-DTT after lesion of ChIs in DMS. Blue dots indicate individual competition time in the first session. Orange dots indicate individual competition time in the last session. Line shows mean of the first or last session in each social rank.

According to our previous results (Table 2A and Figure 3A), there was no significant difference between the control dominants and dominants who won in the between-cage DTT. Therefore, we chose the group of dominants who won in the between-cage DTT as the comparison group. Table 4A shows after the ChI-lesion, dominants who lost in the competition with dominants from other cages did not show a significant decrease in their social rank when tested against their home cage mates. Five of the 8 dominants who lost to other dominants in the between-cage competition nevertheless maintained their rank in the home cage (*p* = 0.2 compared with the comparison group, 2×2 Fisher’s exact test). Only two out of six cages of dominant mice who lost in the between-cage DTT decreased their social ranking relative to cage-mates that had won their between-cage DTT. Furthermore, this loser effect only decreased their ranks compared with the comparison group (day1 – *p* = 0.0288, day2 – *p* = 0.3772, day3 – *p* = 0.3772, day4 – *p* = 0.3772, n = 6, Dunn’s test, Bonferroni adjustment) at the first day of subsequent within-cage DTT (Figure 6A). However, there was no significant difference between unlesioned and lesioned dominants who lost in the between-cage DTT in the context of cage-mates that won during the competition over four consecutive days (Figure 6C, day1 – *p* = 0.5337, day2 – *p* = 0.0734, day3 – *p* = 0.1806, day4 – *p* = 0.0734, Mann-Whitney U test). These results implied that the lesion of ChIs in DMS reduced the loser effect in dominants. Interestingly, there was a significant difference at the third day before the between-cage competition between dominants who lost in the between-cage DTT in the context that their cage-mates lost as well and the comparison group (*p* = 0.0342) or dominants who lost in the between-cage DTT in the context that their cage-mates won in the competition (*p* = 0.0429) (Dunn’s test, Bonferroni adjustment). This result indicated the possibility that the lesion of ChIs in DMS increased the fluctuation of social hierarchy in their home-cage.

**Figure 6.**
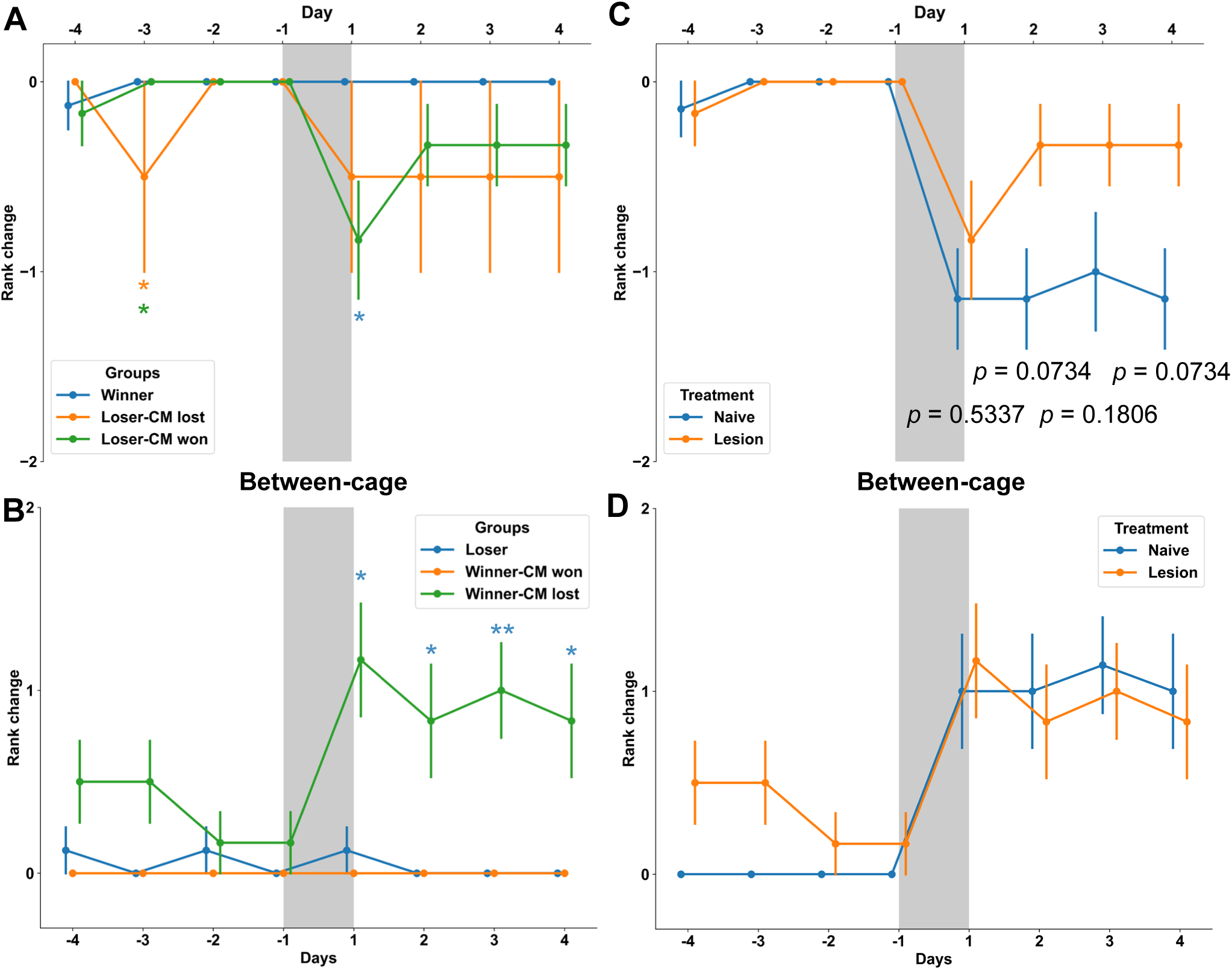
Rank change of subsequent hierarchy after experiencing between-cage competition following lesion of ChIs in DMS. Grey area represents phase of between-cage competition. Horizontal axis indicates the dates before and after between-cage competition. (**A**) Dominants who experienced losing in the context that one or two of their cage-mates won in the between-DTT significantly decreased their ranks in home-cages only at the first day but not at the following days (n = 6) compared with dominants who experienced winning (blue asterisks) in the lesion group (Dunn’s test, Bonferroni adjustment). (**B**) Subordinates who experienced winning in the context that one or two of their cage-mates lost in the between-DTT significantly increased their ranks in home-cages over four consecutive days (n = 6) compared with subordinates who experienced losing (blue asterisks) in the lesion group (Dunn’s test, Bonferroni adjustment). (**C**) In dominants who experienced losing in the context that one or two of their cage-mates won in the between-DTT, there is no significant difference between naïve (n = 7) and lesion group (n = 6) (Mann-Whitney U test). (**D**) In subordinates who experienced winning in the context that one or two of their cage-mates lost in the between-DTT, there is no significant difference between naïve (n = 7) and lesion group (n = 6) (Mann-Whitney U test). (Cage-mates: CM)

**Table 4.** Rank change in home cages after between-cage competition following lesion of ChIs in DMS.

| **A Dominant** |  |  |
| --- | --- | --- |
| Hierarchical changes | Winner in Between-DTT | Loser in Between-DTT |
| Maintained | 8 | 5 |
| Decreased | 0 | 3 |
| Total number | 8 | 8 |

| **B Subordinate** |  |  |
| --- | --- | --- |
| Hierarchical changes | Winner in Between-DTT | Loser in Between-DTT |
| Maintained | 4 | 8 |
| Increased | 4 | 0 |
| Total number | 8 | 8 |

(A) There is no significant difference between dominants who experienced losing and dominants who experienced winning after the lesion of ChIs in DMS ( $p = 0.2$ ). (B) There is also no significant difference between subordinates who experienced winning and subordinates who experienced losing after the lesion of ChIs in DMS ( $p = 0.0769$ ) (2x2 Fisher's exact test).

Regarding the subordinates, there was no significant difference between the control subordinates and subordinates who lost in the between-cage DTT according to our previous results (Table 2B and Figure 3B). Therefore, we chose the group of subordinates who lost in the between-cage DTT as the comparison group. Table 4B shows that subordinates who won the between-cage DTT competition did not show a significant increase in their social ranking in the home cage. Only 4 out of the 8 subordinates who won the between-cage DTT increased their rank in their home cage (*p* = 0.0769 compared with the comparison group, 2×2 Fisher’s exact test). However, after separating the case that their cage-mates also won in the between-cage DTT, the rank of these subordinates (n = 6) showed significant increases compared with the comparison group (day1 – *p* = 0.0213, day2 – *p* = 0.0184, day3 – *p* = 0.0043, day4 – *p* = 0.0184, Dunn’s test, Bonferroni adjustment) over four consecutive days (Figure 6B). There was also no significant difference between unlesioned and lesioned subordinates who lost in the between-cage DTT in the context of cage-mates that won during the competition over four consecutive days (Figure 6D, day1 – *p* = 0.8356, day2 – *p* = 0.8356, day3 – *p* = 0.8356, day4 – *p* = 0.8356, Mann-Whitney U test). These results indicated that the lesion of ChIs in DMS did not alter the winner effect in subordinates.

On the other hand, after lesioning the ChIs a somewhat different pattern of results was seen for intermediate ranked mice who won or lost in the between-cage DTT (Supplementary Table 1). ChI-lesioned intermediates who won in the between-cage DTT tended to maintain their status, resulting in no significant difference from expected outcomes (*p* = 1, 2×3 Fisher’s exact test). Intermediates who lost in the between-cage DTT showed significant differences when compared with expected outcomes (*p* = 0.0256, 2×3 Fisher’s exact test). In the case in which intermediates who won the between-cage DTT had dominant cage-mates that lost in the between-cage DTT (n = 5), there was no significant difference from the expected control (day1 – *p* = 0.8151, day2 – *p* = 1, day3 – *p* = 1, day4 – *p* = 1, Dunn’s test, Bonferroni adjustment) over four consecutive days (Supplementary figure 3). In the case in which intermediates who lost the between-cage DTT had subordinate cage-mates that won in the between-cage DTT (n = 4), there was only significant decreased at the third day of the subsequent within-cage DTT from expected control (day1 – *p* = 0.0914, day2 – *p* = 0.2086, day3 – *p* = 0.0395, day4 – *p* = 0.2086, Dunn’s test, Bonferroni adjustment).

Taken together, our results indicated that the lesion of ChIs in DMS reduced the loser effect in dominants, but did not alter the winner effect in subordinates.

### No association of body weight with social ranking

The possibility that body weight determined social rank (24) was also considered. As shown in supplementary figure 4 and 5, in contrast to the effect of experience of competition with same-ranked mice from other cages, and the experience of cage mates, the body weight of mice was not associated with rank in the social hierarchy in the original home cage in either naïve or lesioned mice.

## Discussion

We investigated flexibility of the transitive hierarchy of male mice, focusing on the effects of experiences of competition outside their home cage, and the role of striatal cholinergic interneurons in these effects. The main findings were as follows. Firstly, position in the transitive hierarchy was flexibly modified by experiences of competition with other mice, external to the home cage. The outcomes of such competition modified the status of mice when they were returned to their home cages. When dominants of one cage lost tube-test competitions to dominants from another, they were subsequently more likely to lose to lower-ranked cage-mates. Conversely, when lower ranked mice won tube-test competitions with similarly ranked mice from other cages, they became more likely to win in competition with higher-ranked mice in their home cage. Secondly, the cholinergic interneurons in the dorsomedial striatum played a selective role in the effect of losing in the external competition on the subsequent hierarchical changes in the home cage in dominants. Thirdly, however, the cholinergic interneurons in the dorsomedial striatum were not involved in the winner effect of the external competition on the subsequent hierarchical changes in the home cage in subordinates. These findings indicate that position in the social hierarchy is dynamic and flexible, and that striatal cholinergic interneurons are involved in experience-related flexibility of the social hierarchy.

We found that the experience of competition with mice of equivalent hierarchical ranking from other cages generated a new social hierarchy when winners and losers returned to their home cages. This finding contrasts with results of an earlier study in which dominant mice that lost to other dominant mice still won competitions against their cage-mates when they returned to their home cages (10). The difference in these results may be because in the present study the cage-mates of the dominants also engaged in competition with mice of similar rank from other cages, and experienced winning or losing. In the earlier study cage-mates did not experience a winning and losing experience against other individuals, so these cage-mates may tend to follow the usual strategy which might cause the social hierarchy to be remained. Our results also supported their findings, for example, the rank reduction only happened in the case that cage-mates won in the contest in dominant-losers, and there was no rank change if their cage-mates lost in the contest as well. According to our results, we suggest that not only the winning or losing experience on target individuals but also the opposite experience on their cage-mates plays a crucial role in the flexibility of social hierarchy.

Our results also showed the correlation between dominance performance in initial and subsequent within-cage WST. This suggests a consistent dominance performance among each cage-mates in home-cage in WST. However, in contrast to previous studies (10, 11, 23) we found a low correlation between DTT and WST. This may be because our cage group size (three mice) was less than the cage group size used previously (four mice). When population density was higher, Brandt’s voles produced more aggression and chasing behavior (25). Thus, a lower population density in a cage may have a lower level of competition among rodents, and which may perform less agonistic behaviors during the WST. In addition, during the analysis of WST, we notice that some groups of mice tended to share the warm spot instead of pushing against the other cage-mates or the competitor. This phenomenon may imply that WST may allow more cooperative behavior than DTT in a lower population density environment.

Previous studies have shown that the prefrontal cortex also contributes to social dominance (10, 11) and behavioral flexibility (26). The involvement of the striatal ChIs in these processes is consistent with the anatomical location and function of the striatum as an entry point for prefrontal input to the basal ganglia (27). Prefrontal inputs trigger dopamine release in dorsal striatum (28), and are important in transmitting prefrontal cortical activity into movement and decision making (29). Although the mechanism by which striatal ChIs contribute to behavioral flexibility is unknown at present, there is evidence that they modulate dopamine-dependent synaptic plasticity in the striatum (30), and respond to unexpected loss (21). Thus, it is possible that striatal ChIs mediate the effects on striatal circuitry of positive and negative outcomes of competition for dominance.

Our study provided new evidence concerning the neural mechanisms that underlie establishing a new social hierarchy, and the winner-loser effect on social hierarchical flexibility. We found that a lesion of ChIs in DMS specifically decreased the loser effects in dominants but not the winner effects in subordinates. This is the first study to indicate that the ChIs in DMS are involved in social hierarchy dynamics and maintenance. Our findings are consistent with previous studies showing that the ChIs in DMS contribute to behavioral flexibility (18–22). In this framework winning or losing is a form of behavioral flexibility which results in a changed strategy. Loss of this flexibility after lesioning ChIs causes difficulty for dominants to switch their strategy from winner to loser in their home-cage. In contrast, lesioning ChIs did not reduce the ability of subordinates to switch from loser to winner.

Behavioral flexibility involves two steps: extinction of the previous cue and learning of the next cue. In relation to extinction, previous studies have shown that ChIs in DMS play an essential role in devaluation and extinction in reversal, but not in the initial devaluation in the reward-based behavioral tasks (19, 20). Thus, it is reasonable to speculate that the activity of ChIs in DMS facilitates extinction during reversal. In this scenario, the status of competitors in the within-cage competition can be considered a context. The dominants tend to use their habitual action (pushing) in the context of intermediate or subordinate competitors and receive the same outcome (winning: a reward). The subordinate tends to use their habitual action (withdrawing) in the context of dominant or intermediate competitors and obtain the opposite outcome (losing: a non-reward). The intermediate, in contrast, actively chooses their action according to the competitors (dominant or subordinate) and obtains the corresponding outcome. On the other hand, learning of a new cue occurs in the between-cage competition. This causes a reversal for one of the same-status competitors, the dominant-losers, and the subordinate-winners. The dominant-losers experience non-reward and thus extinction regulated by the ChIs in DMS, but the subordinate-winner mice mostly experience reward. Therefore, the effect of the lesion of ChIs in DMS could be explained by a reduction of the loser effect in the dominant-loser group, but unchanged winner effect in the subordinate-winner group.

## Materials and Methods

### Animals

Male mice, C57BL/6J (RRID: IMSR_JAX:000664, The Jackson Laboratory) (WT) and heterozygous B6;129S6-Chat^tm2(cre)Lowl/J^ mice (RRID: IMSR_JAX:006410, The Jackson Laboratory) (ChAT-Cre) aged 7 - 16 weeks, were used in the study. Experiments were conducted in accordance with the 2006 guidelines for Proper Conduct of Animal Experiments of the Science Council of Japan and were approved by the Committee for Care and Use of Animals at the Okinawa Institute of Science and Technology, an AAALAC-accredited facility, under protocols (#ACUP-2021-003 and #ACUP-2023-048).

### Behavioral procedures

Mice were housed in groups of three in the home cage for at least two weeks before the start of experimental procedures. A short transparent pipe (15 centimeters length, 3 centimeters diameter) was placed into each cage. During habituation mice were able to naturally establish a social hierarchy that included three social ranks: dominant, intermediate, and subordinate.

To measure the social hierarchy two behavioral tasks were used: the dominance tube test (DTT), a reliable and simple method to determine social ranking in mice (11, 31), and the warm spot test (WST) (11), which was used to provide a comparison with the DTT. The overall timetable is outlined in Figure 1. The DTT apparatus comprised a transparent acrylic pipe (60 cm length, 3 cm diameter) with gates at each end and in the middle. The tube was mounted on a desk with a camera (DFK33UX290, The Imaging Source) to record each trial. Mice were habituated to the apparatus over two consecutive days. During the first day of habituation, mice were individually allowed to freely explore the apparatus. Entries into each end of the pipe were counted, and habituation was terminated when each end of the pipe had been entered at least 5 times. During the second day of habituation, mice were allowed to enter the pipe, but the middle gate was initially closed. The gate behind the mouse (the withdrawal side) was closed after it entered the pipe. After 15 seconds the middle gate was opened to allow the mouse to move forward to the other end of the pipe. The second day of habituation was completed when each mouse entered both ends of the pipe 5 times.

The DTT procedure was initiated by randomly assigning mice to an entry end of the tube. After mice entered the pipe, the three gates were closed. After a 15 second delay, all three gates were opened to allow free competition between the two mice. The trial was completed when one of the mice withdrew from the pipe. This identified the winner and loser of the competition.

The DTT was used to identify the social hierarchy in each home-cage (within-cage DTT). One session was conducted per day. Within-cage DTTs were conducted using all possible pairings of mice in the home cage, to ensure each mouse competed against each cage mate. The within-cage DTT was repeated over 4 to 6 consecutive days to determine the home cage social hierarchy. The hierarchy in the home-cage was considered transitive if the ranking determined by the within-cage DTT remained stable for 4 consecutive days or 5 out of 6 sessions. After each trial the desk and pipe were wiped with 75% ethanol to sanitize the apparatus and remove feces and urine.

The DTT was also used as an experimental treatment inducing competition between mice from different cages (between-cage DTT). To investigate the effects on the home-cage social hierarchy of external competition with mice from other cages, between-cage DTTs were conducted between mice of the same social ranking from different cages.

The WST was used to provide a comparison with the DTT. The WST was used in the same way as the DTT to identify the existing social hierarchy in the home cage (within-cage WST), and to test the effect of external competition on the home-cage social hierarchy (between-cage DTT). There were two stages in the WST: cold adaptation and competition for the warm spot. In the cold adaptation stage, the mice were placed into a plastic box surrounded by ice to cool the environment to around 0 ° C for 30 minutes. The mice were subsequently transferred to another box of the same size to compete for access to a warm platform, a heating pad of 5 centimeters diameter in one corner. The temperature of the heating pad was set at about 33° C. Mice competed for the warm spot for 20 minutes during which their behavior was recorded by a camera for further analysis. The three mice from each home cage were measured together in the within-cage WST, and the same pairs of mice tested in the between-cage DTT were tested together in the between-cage WST.

### Behavioral and statistical analysis

In the DTT, the experimenter measured the duration of competing behavior, from door-opening until one of the mice withdrew, ending the competition. Longer durations in the DTT indicate continued competition, while shorter durations indicate formation of a stabilized hierarchy among competitors. In the WST, the time spent on the warm spot represents the dominance level in each individual mouse. A longer time spent on the warm spot indicates higher dominance in a social group, and vice versa. Videos of the WST were scored using BORIS (32). The first five minutes of the WST was considered habituation, and the time for habituation was excluded from the analysis.

SciPy package in Python was used for statistical analyses. For group comparison of categorical data, we used Fisher’s exact test for small samples. To test the winner and loser effect on rank change over days, we used Dunn’s test with Bonferroni adjustment for post-hoc testing of contrasts. For group comparisons of single continuous variables, we used Student’s unpaired *t*-test. To test session and rank effects and effects of lesions, we used two-way repeated measures ANOVA, with Tukey’s HSD for post-hoc testing of contrasts. To test the lesion effect on rank change over days, we used Mann-Whitney U test for testing contrasts. Pearson’s correlation test was applied to test the correlation between the individual body weight and social rank, and the test was also used to test the correlation between the dominance performance in DTT and WST. No data transformations were used.

### Surgical procedure

Adeno-associated virus (AAV) encoding Cre-dependent diphtheria (AAV1-mCherry-flex-dtA) (33) was used to ablate the ChIs in DMS, and label non Cre-expressing cells with mCherry to confirm the region of viral expression. Mice were anesthetized with 1.5 - 2 % vaporized isoflurane (CYGNI-VET, Japan) during the surgery. Bilateral injections were given of 100 nl virus at two depths (DV: −2.5 and −2.8 mm; AP: 0.7 mm; ML: ± 1.5 mm relative to the bregma) at age 7-10 weeks Carprofen (Rimadyl, 0.5 mg/kg) was subcutaneously injected into the mice after the end of the surgery for pain alleviation. Mice were allowed to recover for at least two weeks.

### Histology

The specificity and extent of the lesion was verified by histological analysis of ChAT-expressing neurons in the DMS after the viral injection. At the end of the experiment mice were deeply anesthetized and then perfused with 4% paraformaldehyde in PBS, and the brains were extracted and postfixed in the same fixative overnight. The brains were then transferred into 30% sucrose in PBS and kept shaking until the brains sank down after which the brains were sectioned on a freezing sliding microtome (REM-710+MC-802A, Yamato) into 30 µm coronal sections. The sections were kept in PBS for further fluorescent immunostaining. To verify the lesion of ChIs and the off-target effect of the viral injection sections were stained for ChAT and NeuN. A 2x HEPES-TSC solution was prepared and modified as previously described (34). This stock solution contained 20% TritonX-100, 20 mM HEPES (4-(2-hydroxyethyl)-1-piperazineethanesulfonic acid), and 400 mM sodium chloride, adjusted to pH of 7.5 by sodium hydroxide. The sections were submerged in blocking solution (1x HEPES-TSC and 5% normal donkey serum) for 1 hour. After blocking, the sections were transferred into primary antibody solution and kept at 4 ° C overnight. The primary antibody solution contained 1x HEPES, 3% normal donkey serum, 1:1000 rabbit anti-NeuN antibody (ABN78, Sigma-Aldrich), and 1:100 goat anti-ChAT antibody (AB144P, Sigma-Aldrich). After three washes in PBS, the sections were transferred into the secondary antibody solution at room temperature for two hours. The secondary antibody solution contained 1x HEPES, 3% normal donkey serum, 1:500 Alexa Fluor 488 donkey anti-rabbit IgG (A21206, Invitrogen), and 1:500 Alexa Fluor 633 donkey anti-goat IgG (A21082, Invitrogen). After a further three washes in PBS, the sections were mounted onto the slides with ProLong Glass Antifade Mountant (P36980, ThermoFisher).

The mounted sections were examined using a confocal (LSM900, Zeiss) or epifluorescence (Dmi8, Leica) microscope. To correct for bleed-through from the emission wavelength of mCherry into Alexa Fluor 633 excited by 640 nm laser (Supplementary figure 5B right), neighboring sections of the same brain were stained without primary antibodies as a negative control (Supplementary figure 5 left). The mCherry bleed-through was corrected by subtraction from the Alexa Fluor 633 signal in the negative control, by adjusting the intensity to obtain parameters that removed the bleed-through (Supplementary figure 5C left). The same parameters were applied into the standard stained image to remove the bleed-through effect on the sections (Supplementary figure 5C right). The corrected image was the representative image shown in figure 3C.

## Acknowledgements

The technical support of the OIST Imaging Section and Animal Resource Sections is gratefully acknowledged. The research was supported by the Okinawa Institute for Science and Technology Graduate University subsidy fund, and HFSP Project Grant PGP0062/2019.

