## Supplementary for "Cholinergic interneurons of the dorsomedial striatum mediate winner-loser effects on social hierarchy dynamics in male mice"

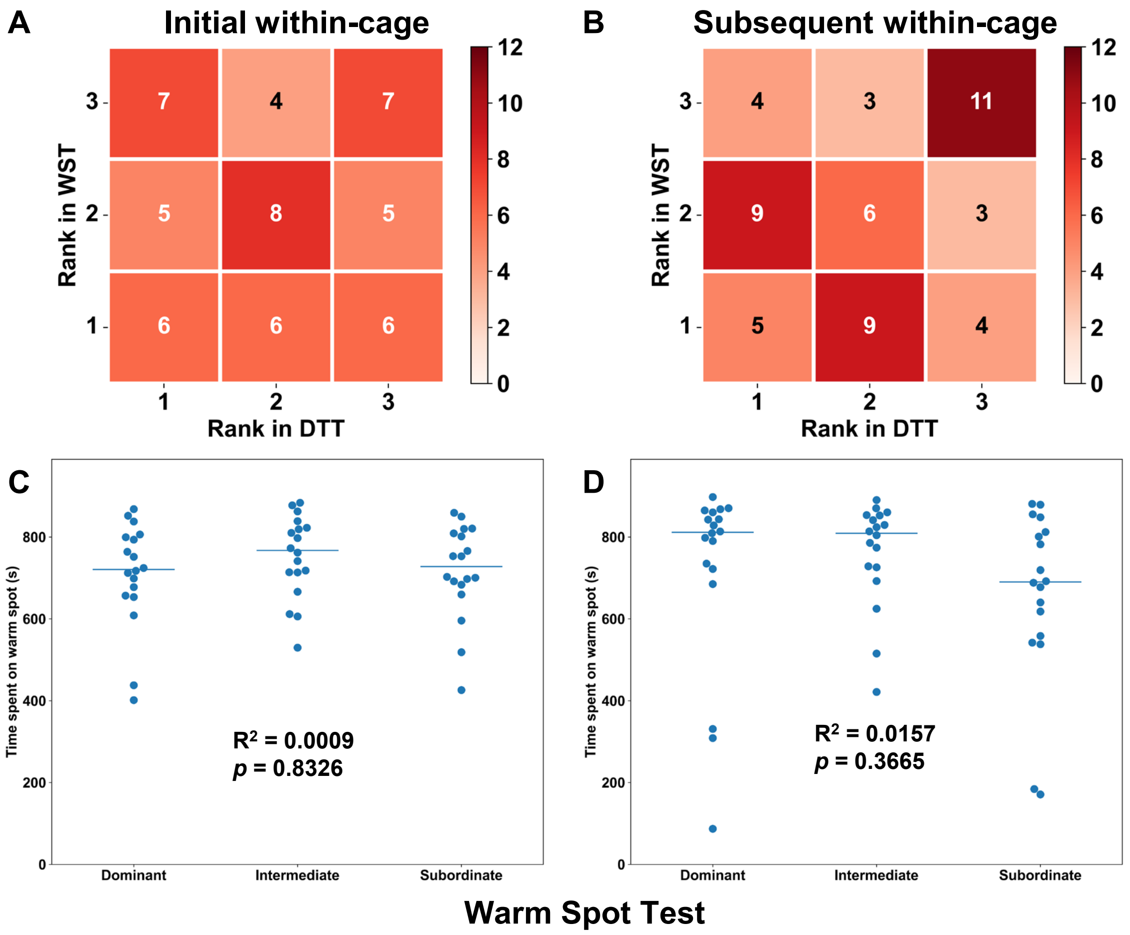


Supplementary figure 1. (**A**) Contingency table with the number of cages shows the correlation between the social hierarchy in DTT and WST in the initial within-cage competition. (**B**) Contingency table with the number of cages shows the correlation between the social hierarchy in DTT and WST in the subsequent within-cage competition. (**C**) The time spent on the warm spot in each social hierarchy in the initial within-cage WST. (**D**) The time spent on the warm spot in each social hierarchy in the subsequent within-cage WST. ($R^2$, *p*, from Pearson's correlation test)

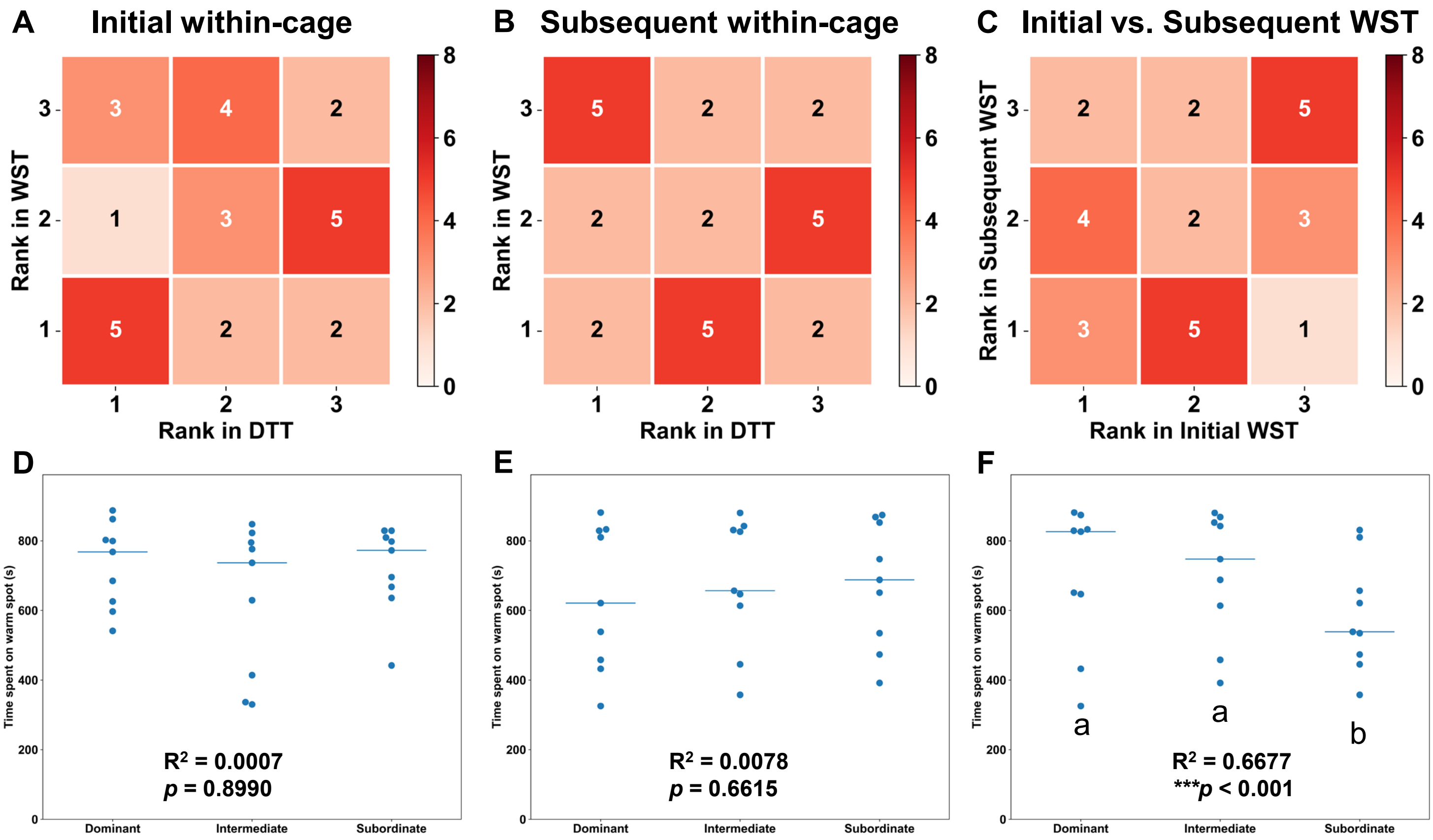


Supplementary figure 2. The contingency tables and time spent on warm spot in control group. (**A**) Contingency table with the number of cages shows the correlation between the social rank in DTT and WST in the initial within-cage competition. (**B**) Contingency table with the number of cages shows the correlation between the social rank in DTT and WST in the subsequent within-cage competition. (**C**) Contingency table with the number of cages shows the correlation between the social rank in initial and subsequent within-cage WST in the subsequent within-cage competition. (**D**) The time spent on the warm spot in each social hierarchy in the initial within-cage WST. (**E**) The time spent on the warm spot in each social hierarchy in the subsequent within-cage WST. (**F**) The time spent on the warm spot in each social rank which was identified by the initial within-cage WST in the subsequent within-cage WST. ($R^2$, *p*, from Pearson's correlation test, Tukey's HSD as the post-hoc test)

Supplementary table 1. Rank change of intermediates in home cages after between-cage competition following lesion of cholinergic interneurons in DMS.

| **Intermediate** | | | |
|---|---|---|---|
| Hierarchical changes | Winner in Between-DTT | Control (expected) | Loser in Between-DTT |
| Increased | 1 | 0 | 1 |
| Maintained | 7 | 8 | 3 |
| Decreased | 0 | 0 | 4 |

Winner vs. Control: $p = 1$; Control vs. Loser: $^*p = 0.0256$

There is no significant difference between intermediates who experienced winning and the expected control. Intermediates who experienced losing significantly changed their social hierarchy in the home cages compared with the expected control. (2x3 Fisher's exact test).

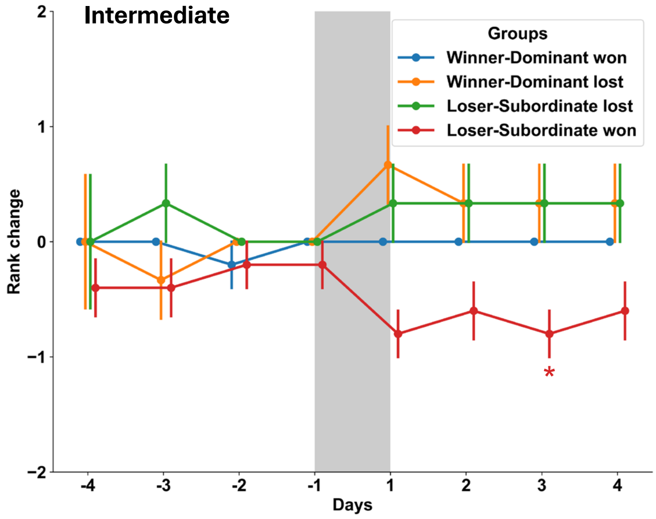


Supplementary figure 3. Rank change of subsequent hierarchy in intermediates after experiencing between-cage competition. Grey area represents phase of between-cage competition. Horizontal axis indicates the dates before and after between-cage competition. Only intermediates who experienced losing in the context that subordinates won in the between-cage DTT significantly decreased their ranks in home-cages at the third day of subsequent DTT (n = 5) compared expected control (red asterisks)). (Dunn's test, Bonferroni adjustment)

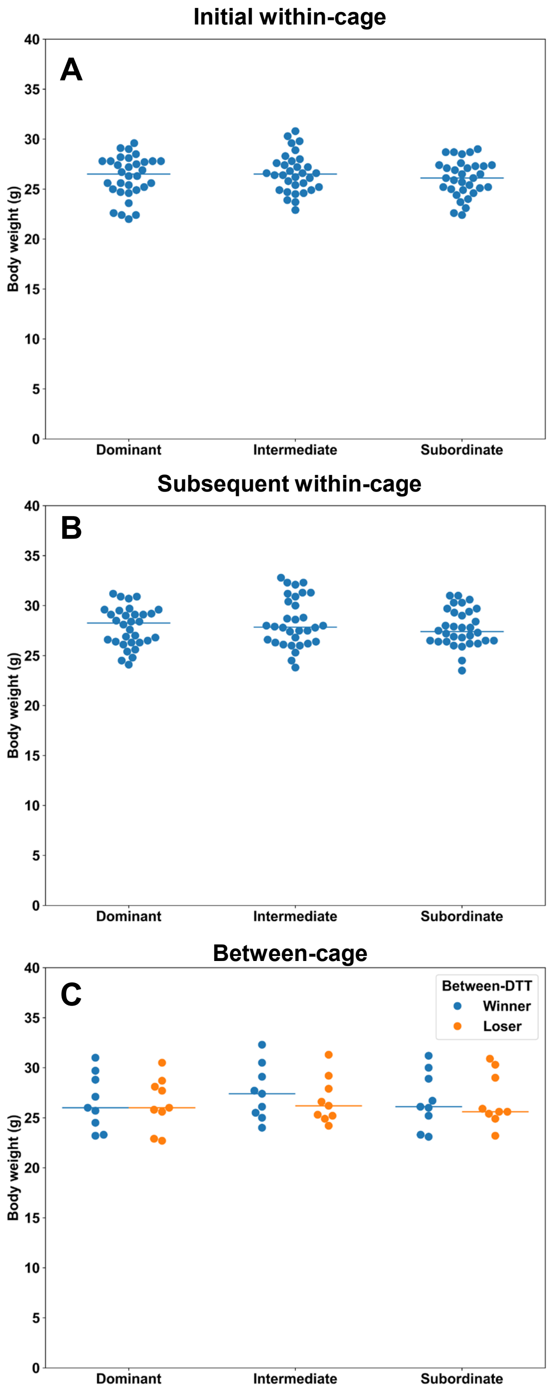


Supplementary figure 4. Body weights of mice in different social ranks. (**A**) Body weight was not significantly different between three social ranks (initial within-cage DTT) ($F_{2,31}$ = 0.032, $p$ = 0.8181, $R^2$ = 0.0001, from Pearson's correlation test). (**B**) Body weight was not significantly different between three social ranks (subsequent within-cage DTT) ($F_{2,31}$ = 0.0231, $p$ = 0.8682, $R^2$ = 0.0005, from Pearson's correlation test). (**C**) Body weight of mice was not significantly different between winner and loser of between-cage DTT in dominant-dominant ($p$ = 0.7929, paired samples $t$ test), intermediate-intermediate ($p$ = 0.2173, paired samples $t$ test), and subordinate-subordinate ($p$ = 0.9433, paired samples $t$ test)

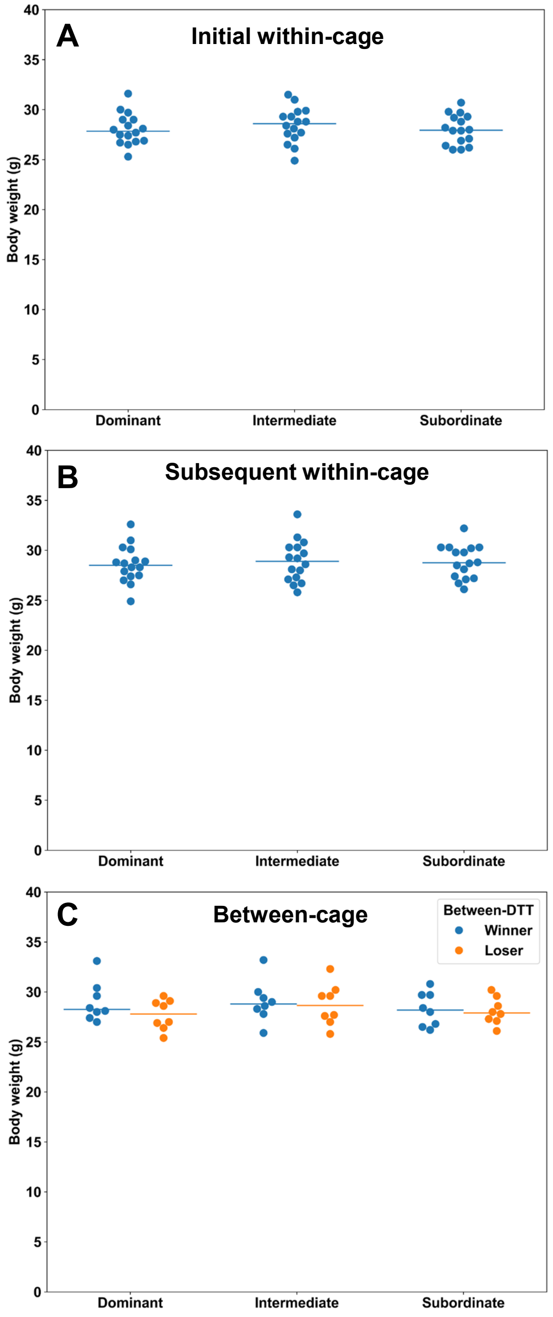


Supplementary figure 5. Body weight of lesioned mice by rank. (**A**) Body weight of lesioned mice was not significantly different between mice of different social ranking determined by the initial within-cage DTT ($F_{2,15}$ = -0.0081, $p$ = 0.9561, $R^2$ = 0.0001, from Pearson's correlation test). (**B**) Body weight of lesioned mice was not significantly different between mice of different social ranking determined by the subsequent within-cage DTT ($F_{2,15}$ = 0.0591, $p$ = 0.6895, $R^2$ = 0.035, from Pearson's correlation test). (C) Body weight of lesioned mice was not significantly different between winner and loser of the between-cage DTT in dominant-dominant ($p$ = 0.1598, paired samples $t$ test), intermediate-intermediate ($p$ = 0.7960, paired samples $t$ test), and subordinate-subordinate ($p$ = 0.8245, paired samples $t$ test).

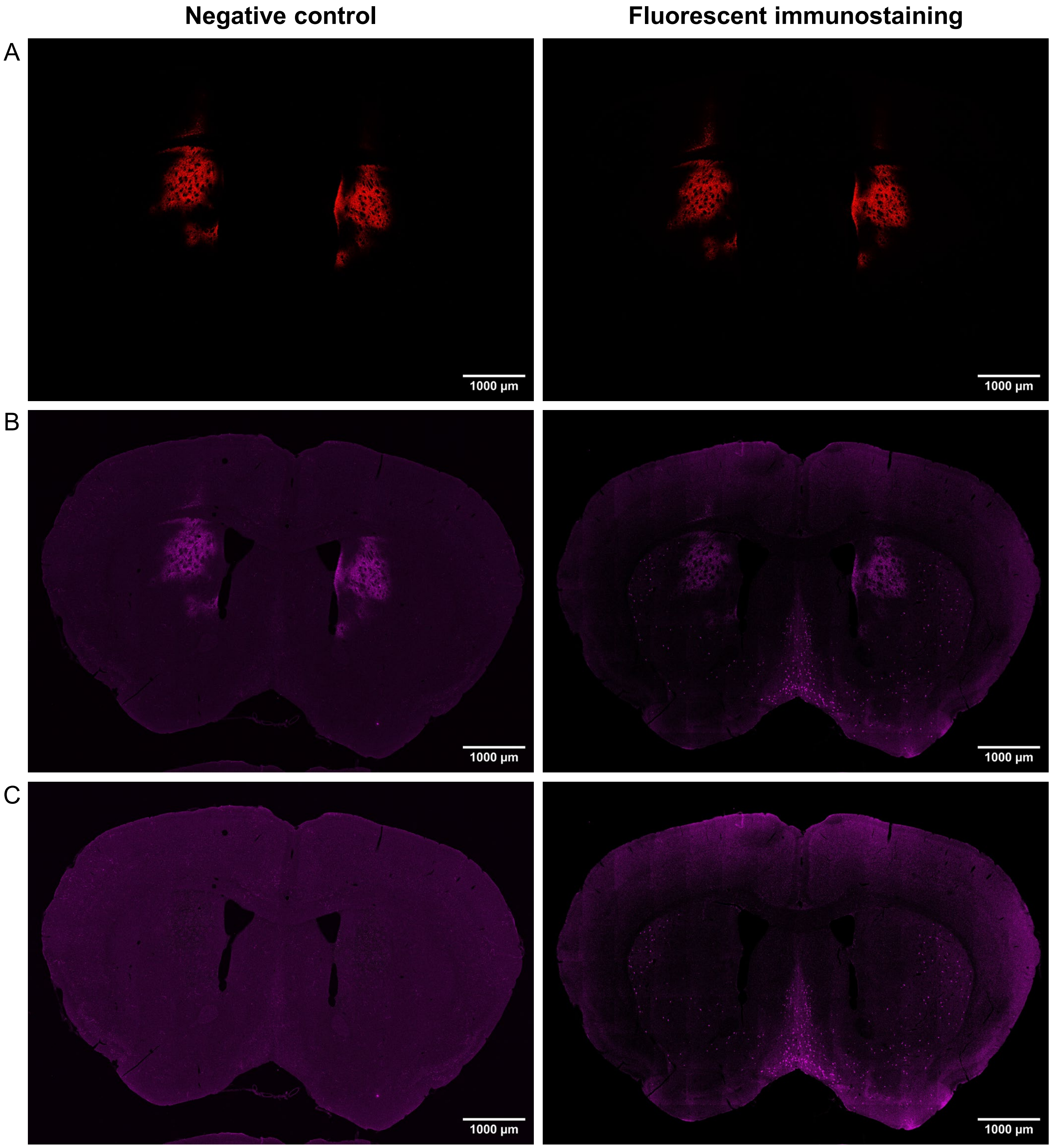


Supplementary figure 6. Process of correcting for bleed-through from mCherry into Alexa fluor 633 emissions when excited by 640 nm laser line. (A) Red shows the mCherry expression in the negative control and immunostaining. (B) The magenta shows bleed-through from mCherry into 650 to 700 nm band in the negative control, which also appears in the right panel using Alexa fluor 633 filter. (C) Imaging of ChAT after subtracting the bleed-through of mCherry in the negative control (left) and ChAT-stained section (right).
